# Proteome-wide analysis of phospho-regulated PDZ domain interactions through phosphomimetic proteomic peptide phage display

**DOI:** 10.1101/211250

**Authors:** Gustav N. Sundell, Roland Arnold, Muhammad Ali, Julien Orts, Peter Güntert, Celestine N. Chi, Ylva Ivarsson

## Abstract

We report phosphomimetic proteomic peptide-phage display, a powerful large-scale method for finding ligands of short linear motif binding domains that simultaneously pinpoint functional Ser/Thr phosphosites in three steps. First, we computationally designed an oligonucleotide library encoding all human C-terminal peptides containing known or predicted Ser/Thr phosphosites and phosphomimetic variants thereof. Second, we incorporated these oligonucleotides into a phage library. Third, we screened the six PDZ (PSD-95/Dlg/ZO-1) domains of Scribble and DLG1 for binding and identified known and novel ligands from the human proteome, and whether these interactions may be regulated by ligand phosphorylation. We demonstrate that the Scribble PDZ domains preferentially bind to ligands with phosphomimetic mutations at two distinct positions, and show that the equilibrium dissociation constant for Scribble PDZ1 with the C-terminal peptide of RPS6KA2 is enhanced over four-fold by phosphorylation. We elucidate the molecular determinants of phosphopeptide binding through NMR structure determination and mutational analysis. Finally, we discuss the role of Ser/Thr phosphorylation as a switching mechanism of PDZ domain interactions.

---

Reversible protein phosphorylation is crucial for regulation of cellular processes, and primarily occurs on Ser, Thr and Tyr residues in eukaryotes ^1^. Phosphorylation may have different functional effects on the target protein, such as inducing conformational changes, altering cellular localization or enabling or disabling interaction sites. Hundreds of thousands of such phosphosites have been identified in different cell lines and under different conditions ^2,3^. An unresolved question is which of these phosphosites are of functional relevance and not background noise caused by the off-target activity of kinases revealed by the high sensitivity in the mass spectrometry analysis. So far, only a minor fraction of identified phosphosites has been linked to functional effects, and only 35% are evolutionary conserved which would support that they play a non-redundant functional roles ^4^. Sifting functional from non-functional phosphosites by experimental approaches is thus of fundamental importance in order to better understand the function of the phosphoproteome.

Phosphosites are often located in the intrinsically disordered regions of the proteins, which in eukaryotes are estimated to cover 30-40% of the populated protein sequence-space ^5^. These regions are enriched in short linear motifs (SLiMs) that are recognized by modular domains. Discrete phosphorylation events may act as a code to regulate and tune the strength of interactions. Phosphopeptide binding domains, such as the 14-3-3 proteins, serve as readers of this phosphorylation code ^6^. Deciphering the phosphorylation code by linking the reader domains to their preferred binding phosphosites is a major challenge. Adding to the complexity, phosphorylation of a SLiM may have a switch-like effect, making interactions stronger (enabling) with a given domain, while weakening interactions (disabling) with other domains^7^, i. e. regulating interactions. Such a switch-like mechanism was recently found to regulate interactions of the C-terminal region of PRTM5 with 14-3-3 proteins and PDZ (PSD-95/Discs-large/ZO-1) domains. The phosphorylation of the C-terminal tail enables 14-3-3 interactions, while disabling PDZ domain interactions ^8^. Although less common, phosphorylation switches also operate on the interaction interfaces of folded proteins, hundreds of which were suggested in recent analysis ^9^.

There is paucity of information linking phosphorylation enabled/disabled SLiMs to their binding partners, in part due to a lack of suitable experimental methods. Although affinity-purification coupled to mass spectrometry (AP-MS) can be used to obtain information on phosphorylation states and interactions dynamically, the approach does not provide information with resolution at the level of the binding sites. In addition, interactions that rely on SLiMs are often elusive to AP-MS as they typically are of low-to-medium affinities, and exhibit rapid association/dissociation kinetics. Phosphopeptide libraries can be used to establish preferred binding motifs of phosphopeptide binding domains, but the number of sequences presented in these experiments are typically limited ^10^. Alternatively, Grossman and co-workers recently showed the use of yeast-two-hybrid (Y2H) for capturing phospho-Tyr-dependent interactions ^11^. Phosphorylation-dependent interactions are otherwise typically not captured by Y2H. Thus, there is an urge for novel large-scale methods for charting phosphorylation-dependent SLiM-based interactions.

Herein, we set out to develop a novel large-scale approach termed phosphomimetic proteomic peptide phage display (ProP-PD) to identify interactions that are enabled and/or disabled by Ser/Thr phosphorylation by combining computational library design, phosphomimetic mutations (Ser/Thr to Glu), custom oligonucleotide library, phage display, and next-generation sequencing (NGS). The phosphomimetic ProP-PD is a further development of ProP-PD, which is an emerging method for the identification of SLiM-based interactions on large-scale ^12^. In a previous study we created a phage library that displays all C-terminal regions of the human proteome (50,549 heptamer peptides). This library was used in selections against a set of PDZ (PSD-95/Dlg/ZO-1) domains, which are among the most frequent interaction modules in eukaryote proteomes with about 267 instances in over human 150 proteins ^13,14^. Binding-enriched phage pools were analyzed by NGS, which provided information on the binding peptides from which the full-length proteins could be identified ^12^,^15^. In addition, the analysis provided an affinity ranked list of ligands as the differences observed in NGS counts for peptides identified within a given selection typically translate into affinity differences. More recently, we created ProP-PD libraries covering a large part of the disordered regions of the human proteome ^16^, and used it to elucidate the substrate recognition of the protein phosphatase 2A B’ holoenzymes ^17^. ProP-PD is thus a well-established method for interaction profiling but, in its original design, could not be used for the identification of phosphorylation switches.

We here show that phosphomimetic ProP-PD is a straightforward approach for finding potential ligands of peptide-binding domains and for simultaneously pinpointing Ser/Thr phosphorylation events with potential to enable or disable interactions. We showcase the approach by identifying phospho-regulated interactions of PDZ domains). PDZ domains are well-known for binding to C-terminal peptides with typical PDZ binding motifs (PDZbms; class I binding motif S/T-x-Φ-coo-, class II Φ -x-Φ-coo-, class III D/E -x-Φ-coo-, where x is any amino acid and Φ is a hydrophobic amino acid) but also interact with internal PDZbms and phospholipids ^14^. The last amino acid in the PDZbm is denoted p 0, and the upstream residue is denoted p -1, and so on. Phosphorylation of the Ser/Thr at p -2 of the class I PDZbm typically disables PDZ interactions ^18–23^, but phosphorylation of other positions may either enable or disable interactions ^24^. First, we created a phosphomimetic ProP-PD library displaying all C-terminal regions of the human proteome containing known or putative phosphorylation sites, and the phosphomometic mutants of these peptides. Second, we used this library in selections against the six PDZ domains of human Scribble and DLG1, well-characterized PDZ proteins with crucial roles in the postsynaptic density of excitatory neuronal synapses and in the establishment and maintenance of epithelial cell polarity ^24,25^. Third, we successfully identified known and novel interactions, with potential to be regulated by Ser/Thr phosphorylation, and uncover that Scribble PDZ interactions are enabled by p-3 phosphorylation. Finally, we provide the structural basis of such phosphopeptide binding and discuss the role of Ser/Thr phosphorylation as a switching mechanism of PDZ domain interactions.

In this proof-of-concept study we thus demonstrate that phosphomimetic Pro-PD is a powerful approach to identify SLiM-based interactions in the human proteome that are regulated by Ser/Thr phosphorylation. The approach is readily scalable and can be used to explore the potential phospho-regulation of human protein-protein interactions on a large scale.

## Results

### Construction of a phosphomimetic ProP-PD library

To design a specific phage display library for phosphomimetic ProP-PD we scanned the human proteome for C-terminal regions containing known or predicted Ser/Thr phosphorylation sites based on phosphorylation databases and *in silico* predictions (see on-line Methods). We identified 4,828 unique C-terminal peptides of human full-length proteins that contain one or multiple potential Ser/Thr phosphosites in their last nine amino acids. We designed the phosphomimetic ProP-PD library to comprehensively contain wild-type sequences and phosphomimetic mutants (Ser/Thr to Glu) thereof (7,626 phosphomimetic sequences, including combinations of multiple phosphorylation sites in one C-terminus), in order to allow the detection of interactions that are enabled or disabled by phosphomimetic mutations. The library design contains 24% of all C all non-redundant human C-terminal peptides of human full-length proteins reported in the annotated section of Swissprot/Uniprot. These sequences conatin 8.2, 6.2 and 3.4% class I, II and II PDZ binding motifs, respectively. In case of class I containing peptides, 40.8% of the phosphorylation sites are at p-2, and are expected to disable PDZbm-dependent interactions.

Oligonucleotides coding for the wild-type and phosphomimetic (Ser/Thr to Glu) peptides flanked by annealing sequences were obtained as a custom oligonucleotide library and incorporated into a phagemide vector, thereby creating a C-terminal fusion proteins of the M13 major coat protein p8, previously engineered for C-terminal display ^26^ (Fig. 1). We confirmed 89% of the designed sequences by NGS and the vast majority of reads matched the library design. Single nucleotide substitutions were present in 6% of the sequences of which 75% are nonsynonymous. The library coverage and quality was thus considered satisfactory.

**Figure 1.**
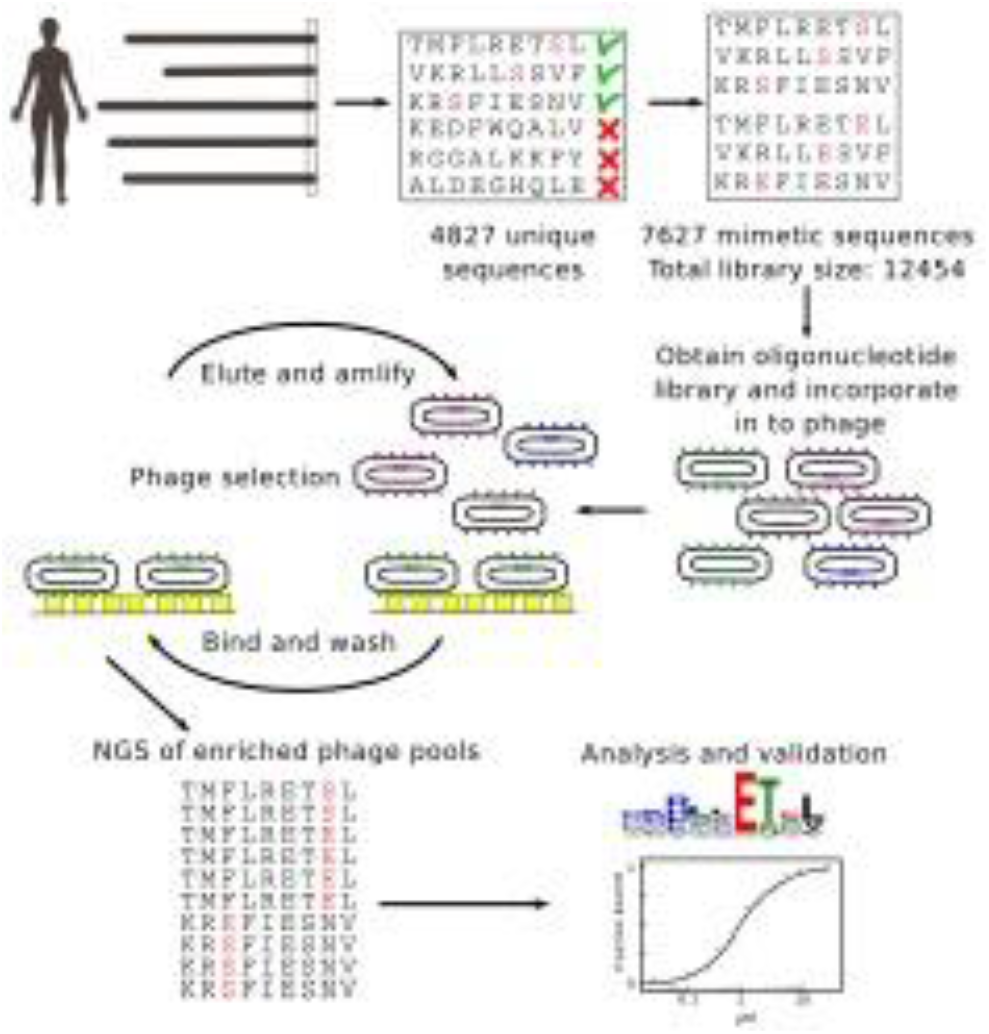
Schematics of the phosphomimetic ProP-PD approach and validations. A phosphomimetic ProP-PD library was designed to display the C-terminal regions (nine amino acids) of all human proteins containing known or putative Ser/Thr phosphorylation sites (4,827 peptides), and all phosphomimetic (Ser/Thr to Glu) variants thereof (7,626 peptide). Oligonucleotides encoding the sequences were incorporated into a phagemid designed for the display of peptides fused to the C-terminus of the major coat protein P8 of the M13 filamentous phage. The library was used in binding selections against immobilized PDZ domains of Scribble and DLG1. Binding enriched phage pools were analyzed by NGS, followed by data analysis and validations.

### Phosphomimetic ProP-PD selections

We tested the performance of the phosphomimetic ProP-PD library through selections against the immobilized PDZ domains of human Scribble (PDZ1, PDZ2 and PDZ3) and DLG1 (PDZ1, PDZ2 and PDZ3). Of these proteins, Scribble has previously been shown to interact with the p-1 phosphorylated PDZbm of MCC (colorectal mutant cancer protein; RPHTNETpSL-coo-)^27^. In contrast, phosphorylation of the p-2 position has been shown to disable interaction with DLG1 PDZ2 ^28^. The experiment was thus designed to capture interactions that were enabled or disabled by phosphomimetic mutations. The selections were successful as judged by phage pool enzyme-linked immunosorbent assay (ELISA) and saturated after three rounds of selection. Binding-enriched phage pools were barcoded and analyzed by NGS, which resulted in a list of binding peptides ranked by their occurrence in the NGS results. High-confidence sets of ligands (Table S2) were obtained by assigning cut-off values that filtered out non-specifically retained peptides lacking typical PDZbms. For each domain, we generated position weight matrices (PWM) based on the ProP-PD (Fig. 2a). All domains are class I binding PDZ domains, and the PWMs are in good agreement with previous studies ^12,29,30^.

**Figure 2.**
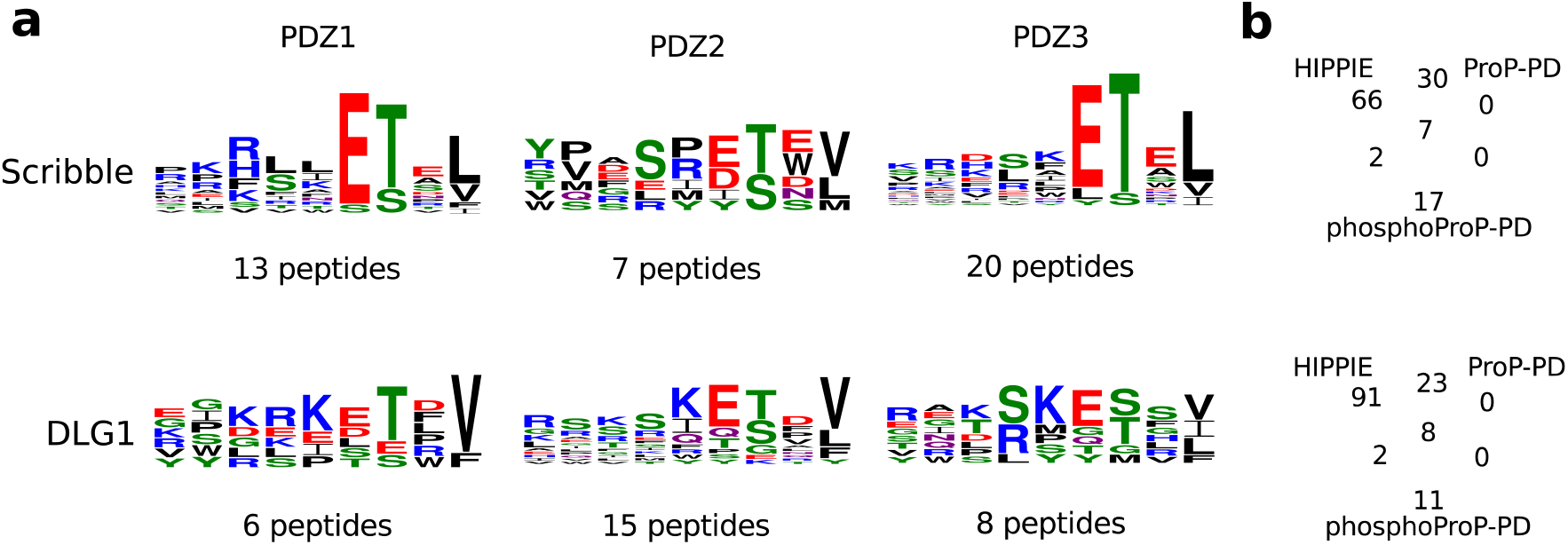
Analysis of phosphomimetic ProP-PD selection data. **a.** Position weight matrices (PWMs, web logo 3) representing the phosphomimetic ProP-PD selections data for the PDZ domains of Scribble and DLG1. The numbers of peptides used to create the PWMs are indicated. **b.** Overlap between ligands identified through phosphomimetic ProP-PD and interactions reported in the HIPPIE database. The HIPPIE database already contains ligands previously reported through ProP-PD ^12^ for Scribble and DLG1, and the overlaps between the two studies are indicated.

### Comparison between phosphomimetic ProP-PD results and known ligands

We compared the ligands identified through phosphomimetic ProP-PD with the known interactions listed in the HIPPIE database as of August 2017 ^31^ (Fig. 2b). HIPPIE incorporates information from databases such as IntAct^32^ and Biogrid ^33^, and contains already our previously generated ProP-PD results for the PDZ domains of Scribble and DLG1. Of the putative Scribble interacting proteins identified in this study, 35% are reported as Scribble ligands in the HIPPIE database, of which 78% have evidence from our earlier study. Among the known interactions found through phosphomimetic ProP-PD, we note the interaction between Scribble PDZ3 and the Hippo pathway protein WWTR1 (KSEPFLTWL-coo-), which, alongside with YAP1, functions as a transcriptional co-activators downstream of the Hippo pathway ^34^. Interestingly, our results also suggest that Scribble PDZ3 interacts with YAP1 (DKESFLTWL-coo-), which provides an additional link between the epithelial cell polarity and the Hippo signaling pathway. Both of these interactions are disabled by phosphomimetic mutations of sites upstream of the core PDZbms. In addition to the ligands reported in HIPPIE, two of the interactions were previously predicted and validated by Luck et al ^13^. For DLG1, 48% of the ligands identified through phosphomimetic ProP-PD have previously been reported by other studies, of which 80% were also found in our pilot ProP-PD study.

The overlap between ProP-PD and phosphomimetic ProP-PD demonstrate that the method robustly reports on overlapping sets of ligands despite differences in library designs such as different peptide length. The previously reported ligands are dominated by interactions that appear disabled by phosphomimetic mutations (78% for Scribble, 90% for DLG1). In contrast, the novel ligands are dominated by interactions that are enabled by the phosphomimetic mutations (78% of novel Scribble PDZ ligands, 55% of DLG1 PDZ ligands). Thus, phosphomimetic ProP-PD allows the identification of novel interactions potentially enabled by phosphorylation as discussed below, thereby complementing protein-protein (PPI) data sets generated by other methods.

### Analysis of the effects of phosphomimetic mutations

We systematically evaluated the effects of phosphomimetic mutations at distinct positions of the PDZbms by analyzing the NGS counts received for each peptide pair (wild-type and phosphomimetic; Fig. 3a-b). Essentially, we calculated the ratio between the NGS counts of a given phosphomimetic peptide and the sum of NGS counts of the phosphomimetic peptide and its wild-type (Table S2). We then calculated the average ratios for each position of the PDZbm. This resulted in a score in the range between 0 and 1 for each peptide position, where 0 indicates that the selection was dominated by wild-type peptides and suggests that phosphomimetic mutations at this position disable interaction. A score of 1 suggests that phosphomimetic mutations at the given position enable interactions. The scores for the different domains and peptide positions are summarized in matrices (Fig. 3b), which reveals common and distinct features.

**Figure 3.**
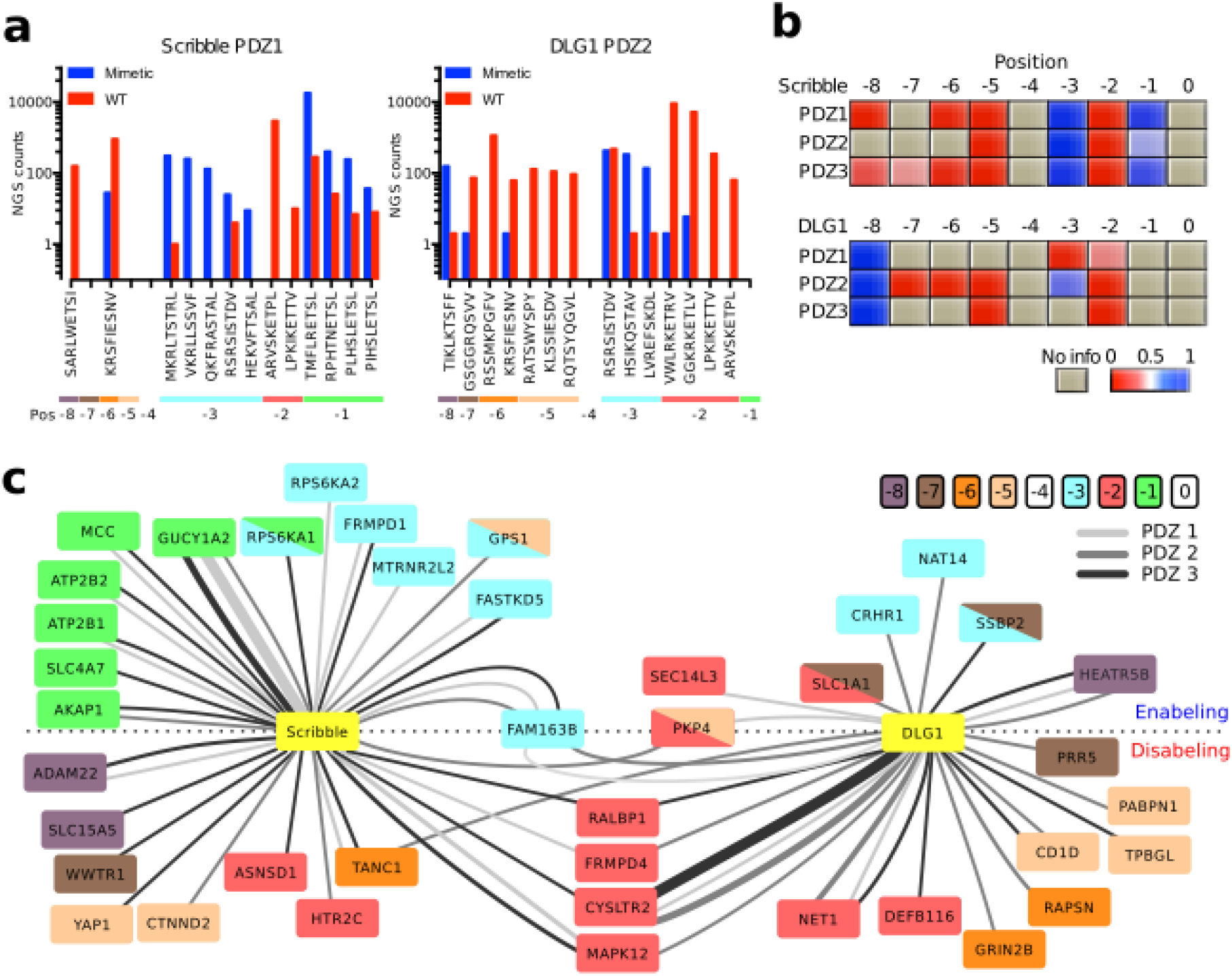
Phosphomimetic ProP-PD reveals peptide position specific effects of phosphomimetic mutations. **a.** Bar graphs of representative NGS data of phage pools enriched for binders through selections against Scribble PDZ1 and DLG1 PDZ2. Peptide sequences are sorted based on the positions of their known or putative phosphorylation sites. Red bars represent WT reads and blue mimetic reads. **b.** Scoring matrices of the effects of phosphomimetic mutations on the phosphomimetic ProP-PD results of the PDZ domains of DLG1 and Scribble. Scores are calculated from the ratios of NGS counts of the phosphomimetic peptides and the sum of the NGS counts of the wild-type and the phosphomimetic peptides. A score of 0 (red) indicates that the selection was dominated by wild-type peptides and a score of 1 (blue) indicates that the selection was dominated by peptides with phosphomimetic mutations at the given position. **c.** Network representation of the putative protein-protein interactions suggested through phosphomimetic ProP-PD. The node color indicates the position of the known or predicted phosphorylation site. Double mutants are indicated by two colors within one node. The edge color indicates the domain (PDZ1, PDZ2, PDZ3) that was found to interacts with the target. The edge thickness reflects the total count of the peptide (WT and phosphomimetic) in the NGS analysis. For targets above the dotted line, the phosphomimetic mutations enable interactions. For targets below the dotted line, interactions are disabled by mutations. Interactions with targets on the dotted line are enabled or disabled depending on which PDZ domain they interact with.

A shared feature is that phosphomimetic mutations at p-2 disable interactions with all PDZ domains, as expected for class I binding PDZ domains. A distinct feature is that phosphomimetic mutations at p-1 enable interactions with Scribble PDZ domains (scores in the range of 0.67-0.93), but not with the DLG1 PDZ domains. Phosphomimetic mutations at p-3 position enable interactions with Scribble PDZ domains and to less extent with DLG1 PDZ2. Phosphomimetic mutations upstream of the core PDZbm disable interactions, with the exception for the TIKLKTSFF-coo- peptide of HEATR5B, that appears to bind better to the DLG1 PDZ domains when Thr at p-8 is replaced with Glu; this may however be an artefact as the notion is based on the observation of only one peptide sequence and residues so far upstream of the PDZbm typically have less impact on the affinity.

We visualized the multilevel information obtained through phosphomimetic ProP-PD in a PPI network to provide an overview of the results (Fig. 3c). In this network, we indicate for which PDZ domain the interaction was identified (edge color), if the phosphomimetic ProP-PD results suggest the interaction to be enabled or disabled by phosphomimetic mutation (above/below dotted line, respectively), and the position of the known or putative phosphorylation site (node color). As can be seen from the network, several proteins are identified as ligands for more than one PDZ domain. From the network representation, it is also obviouss that there are two ligand positions where phosphomimetic mutations appear to enable interactions, namely p-1 and p-3 and in particular for interactions with the PDZ domain of Scribble. In addition, it is noteworthy that phosphomimetic mutation may have a switching function, such that it enables interactions with one protein, while disabling interactions with the other protein, as seen for the p-3 mutation of FAM163B.

### Analysis from the perspective of the kinases

An added value of the phosphomimetic ProP-PD analysis is that the information on the kinases involved in phosphorylating their ligands is included in the library design when available. We therefore evaluated the phosphomimetic ProP-PD data from the perspective of the kinase that are known or predicted to phosphorylate the identified ligands (Table S1). From this analysis, we found that Scribble PDZ domains interact with potential PKA and RSK substrates more frequently than expected based on the frequencies of these kinases’ substrates in the library design (P-values 0.0372 and 0.0372 for PKA and RSK kinases, respectively, Fisher exact test, BH correction for multiple testing), and these phosphorylation events are in all but one case expected to enable the interactions with Scribble. Interestingly, three of the identified Scribble ligands are kinases (RPS6KA1, RPS6KA2, MAPK12). The PDZ domains of DLG1 instead preferentially binds to PKG and CAM-II substrates (P-values 0.0009, 0.005, Fisher exact test, BH correction for multiple testing): Indeed, the DLG1 PDZ domains enriched two out of the only five available CAM-II substrates in the library design. In these cases, phosphorylation is expected to have negative effects on the interactions as the phosphomimetic mutations of the residues confer loss of interactions. The localization of DLG1 itself is proposed to be regulated by CAMK2D and both have been found in complex ^35^.

### Microscale thermophoresis affinity mesurements confirm phosphorylation switches

To explore the extent to which the phosphomimetic ProP-PD results translats into affinity differences between unphosphorylated and phosphorylated peptides we focused on Scribble PDZ1 ligands. We selected three distinct sets of peptides (unphosphorylated, phosphomimetic mutant and phosphorylated) for affinity determination by microscale thermophoresis (MST; Fig. 4a). The peptides were chosen in order to validate the effects of the putative phosphorylation switches at the p-1 (MCC; HTNETSL-coo-), p-3 (RPS6KA2, RLTSTRL-coo-) and the upstream p-6 position (TANC1, KRSFIESNV-coo-). For these peptides, phosphomimetic ProP-PD resulted in a 15-fold enrichment of the phosphomimetic MCC peptide over the wild-type peptide, a 300-fold enrichment for the phosphomimetic RPS6KA2 peptide over its wild-type, and a 32-fold enrichment of the wild-type over the phosphomimetic peptide for the TANC1 peptide. Scribble PDZ1 was titrated against FITC-labeled wild-type, phosphomimetic and phosphorylated peptides. All peptides bound to Scribble PDZ1 with micromolar affinities (0.6-101 µM K_D_ values, Fig. 4a). Consistent with the phosphomimetic ProP-PD results Scribble PDZ1 has a slightly higher affinity for phosphorylated MCC and significantly higher affinity for phosphorylated RPS6KA2, as compared to their unphosphorylated counterparts. The opposite is true for the TANC1 peptide (Fig. 4a). Thus, there is a good qualitative agreement between the phosphomimetic ProP-PD data and the affinity differences between phosphorylated and unphosphorylated ligands as determined by MST for this domain.

**Figure 4.**
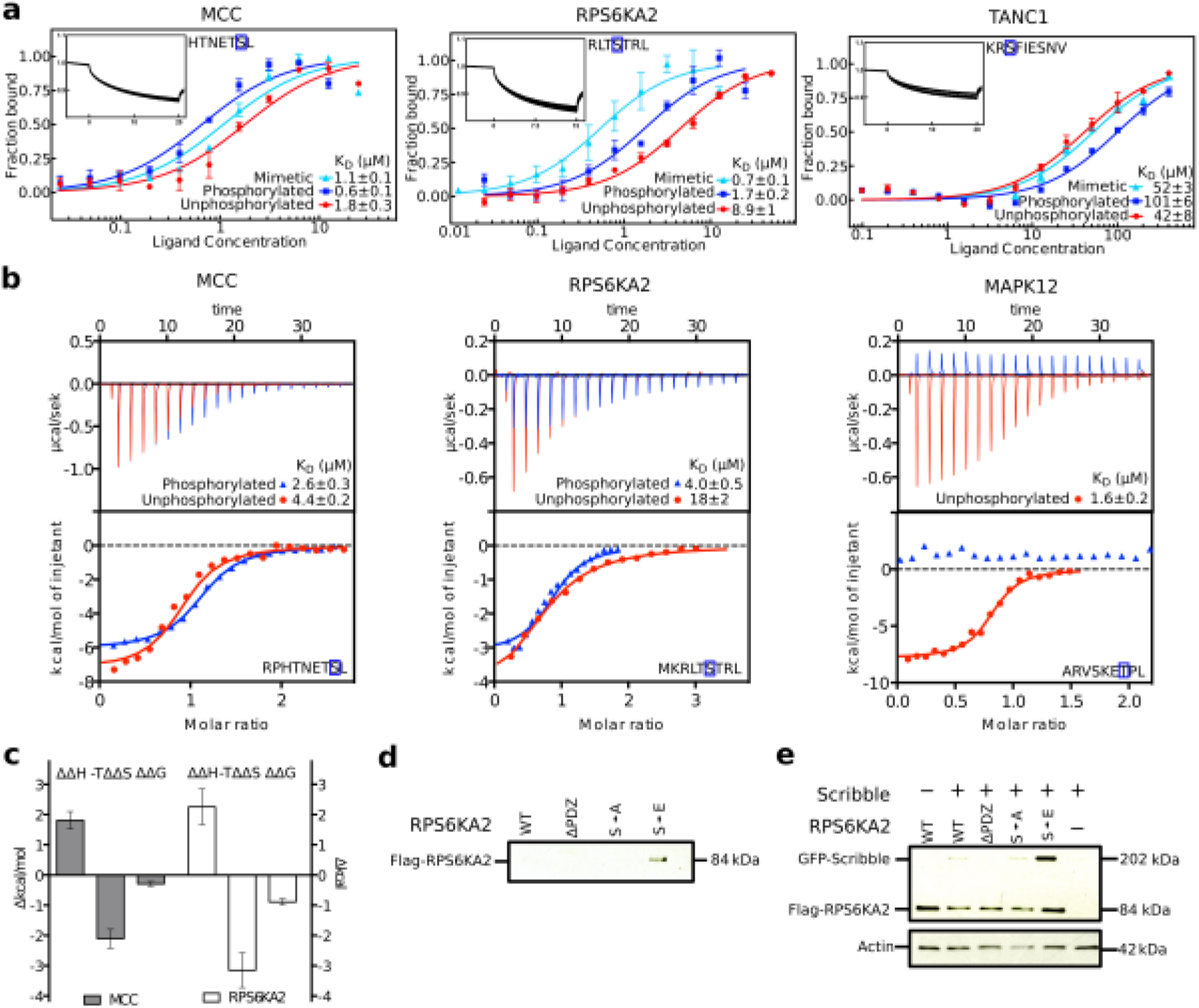
Scribble PDZ1 preferentially interacts with p-1 and p-3 phosphorylated ligands as shown by affinity determinations (MST and ITC) and co-immunoprecipitations. **a.** MST affinity measurements using FITC-labeled peptides (unphosphorlated, phosphorylated and phosphomimetic) of MCC (p-1), RPS6KA2 (p-3) and TANC1 (p-7). A fixed peptide concentration (25–50 μM) was titrated with varying concentration of Scribble PDZ1. *K*_*D*_ values were determined using thermophoresis and T-Jump signal for data analysis. (*n=3*; error bars represent SD). Insets show representative titrations **b.** Representative ITC titrations of unphosphorylated and phosphorylated peptides of MCC, RPS6KA2 and MAPK12 and Scribble PDZ1. The peptides were titrated in to a constant concentration of Scribble PDZ1 (*n=3*). **c.** Thermodynamic parameters of Scribble PDZ1 binding unphosphorylated and phosphorylated MCC and RPS6KA2 as determined in b). **d.** Phosphomimetic mutation of RPS6KA2 (S730E) is required for interaction with Scribble as shown by GST-pulldown. GST-pulldown of full-length Flag-tagged RPS6KA2 WT, mutants (S730A and S730E) and truncated (S730∆) over-expressed in HEK 293T cells with GST-tagged Scribble PDZ1 (left). Detection was performed using an anti-Flag antibody **e.** Co-IPs of GFP-tagged full-length Scribble with Flag-tagged full-length RPS6KA2 constructs as indicated. Detection was performed using an anti-Flag antibody and an anti-GFP antibody.

To evaluate the effect of phosphorylation on DLG1 PDZ binding, we determined the affinities for the same set of peptides with DLG1 PDZ2 (Fig S1) and found them to be in the µM range (2.7-95 µM). Consistent with the indications from the phosphomimetic ProP-PD data (Fig. 3a), phosphomimetic mutation of p-3 improved the affinity of DLG1 PDZ2 for RPS6KA2 significantly (7x). However, phosphorylation of p-3 has only a minor enabling effect on the interaction (1.3x), suggesting that care needs to be taken on a case-to-case basis when evaluating phosphomimetic ProP-PD results. Both phosphorylation and phosphomimetic mutation of the p-6 position of the TANC1 peptide have minor disabling effects (2.9x and 2.4x, respectively). We did not identify any DLG1 PDZ2 ligands with known or putative phosphosites at the p-1 position through phosphomimetic ProP-PD, but MST measurements with p-1 phosphorylated MCC peptide revealed that phosphorylation of this site has a disabling effect on DLG1 PDZ2 interactions (9x). Thus, ligand phosphorylation has differential effects on interactions with distinct PDZ domains, and may serve to switch ligand interactions between different PDZ domains.

### Isothermal titration calorimetry uncovers enthropic-enthalpic compensation

To further confirm the affinity differences determined for Scribble PDZ1, and to gain insights into the thermodynamics of the phosphopeptide interactions we performed isothermal calorimetric (ITC) affinity measurements of Scribble PDZ1 binding to the unphosphorylated and phosphorylated peptides of MCC (p-1 RPHTNETSL-coo-) and RPS6KA2 (p-3 MKRLTSTRL-coo-) (Fig. 4b). In addition, we tested the effect of p-2 phosphorylation using unphosphorylated and phosphorylated MAPK12 peptides (ARVSKETPL-coo-). Consistent with the phosphomimetic ProP-PD and MST results, phosphorylation of MCC at p-1 has a minor enabling effect (2x lower *K_D_)* on the interactions, and phosphorylation of RPS6KA2 p-3 has an enabling effect (5x). Phosphorylation at p-2 of MAPK12 disables the interaction, to the extent that no interaction was observed within the concentration range used (25-2200 µM), in line with the disabling effect of p-2 phosphorylation suggested by the phage display.

The differences in Gibbs free energy of binding (*ΔΔG*^*phos*^ = *ΔG*^*phos*^ *−ΔG*^*unphos*^) between phosphorylated and unphosphorylated peptides were modest, -0.29 kcal/mol for MCC and −0.89 kcal/mol for RPS6KA2 (Fig. 4c). We note an entropy-enthalpy compensation effect for the unphosphorylated-phosphorylated peptide pairs, such that although the interactions were largely enthalpy driven for the unphosphorylated peptides the binding to the phosphorylated peptides were entropy driven. This effect is particularly noticeable for the RPS6KA2 peptide pair, where the change in the entropic contribution to binding upon phosphorylation (*TΔΔS=-TΔSp*^*hos*^*-(-TΔS*^*unphos*^)) is −3.2 kcal/mol (Fig. 4c). Taken together, affinity measurements through MST and ITC confirm that phosphomimetic ProP-PD can be used to identify interactions and to pinpoint putative phosphorylation switches, and the thermodynamic parameters reveal that the effects of phosphorylation on ligand binding is more complex than revealed by the modest affinity changes.

### GST-pulldown and co-immunoprecipitation of Scribble and RSK6A2

We next aimed to verify if the identified interactions and putative phosphorylation switches are functional in the context of the full-length proteins. We focused on the interaction between Scribble and RPS6KA2 as MCC and Scribble already have been shown to interact in a p-1 phosphorylation-dependent manner ^27^. We performed GST-pulldown experiments between GST-tagged Scribble PDZ1 and Flag-tagged RPS6KA2 wild-type (WT), phosphomimetic mutant (S730E), Ala mutant (S730A), and a deletion construct where the three last amino acids had been truncated (RPS6KA2Δ). We successfully confirmed an interaction with RPS6KA2 S730E, but not with the other constructs, supporting that the phosphomimetic mutation is required for interaction (Fig. 4d). In addition, we performed co-immunprecipitations (IPs) of transiently expressed GFP-tagged full-length Scribble and the Flag-tagged RPS6KA2 constructs, and confirmed that full-length Scribble only shows co-IP with the phosphomimetic RPS6KA2 S730E. The results thus support that phosphomimetic ProP-PD can identify putative phosphorylation-dependent interactions that are functional in the context of the full-length proteins.

However, there is so far no experimental support for RPS6KA2 S730 phosphorylation.v In contrast, the corresponding S732 at the p-3 position in the C-terminal tail (VRKLPSTTL-coo-) of the homologous RPS6KA1 is a confirmed phosphorylation site ^36^. We also found that Scribble bound to the phosphomimetic VRKLPETTL-coo- peptide of RPS6KA1 in phosphomimetic ProP-PD (Fig. 2, Table S1). We therefore determined the affinities of unphosphorylated and phosphorylated RPS6KA1 peptide for Scribble PDZ1 through MST using FITC-labelled peptides, resulting in a *K*_*D*_ of 0.39 µM for the phosphorylated RPS6KA1 and 1 µM for the unphosphorylated peptide (Fig. S1, Table S3). The results thus confirm that Scribble PDZ1 preferentially interacts with phosphorylated RPS6KA family members.

### NMR structural analysis of Scribble PDZ1 interactions with unphosphorylated and phosphorylated ligands

To further explore the novel interactions that we identified by phosphomimetic ProP-PD and confirmed through MST, ITC and co-IP experiments, we determined the NMR structure of the Scribble PDZ1 domain when bound to the phosphorylated RPS6KA2 (p-3) peptide. We also performed ^15^N-^1^H heteronuclear single quantum correlation spectrum (HSQC) titrations of the RPS6KA2 and MCC peptides (phosphorylated and unphosphorylated). The bound Scribble PDZ1 structure has the typical PDZ fold, containing of six β-strands and two α-helices arranged in a sandwich pattern (Fig. 5a). The structure was overall similar to the free protein except for the noticeable opened binding pocket and the presence of an extended C-terminal region comprising residues 816-829, which is packed onto the β2/β3 loop. This extended C-terminal region seems to suggest a role of extra structural elements out-side the canonical PDZ-fold. This region was not determined in the free Scribble PDZ1 domain, perhaps because the sequence used for the structure determination was shortened both at the C- and N-termini (PDB codes 1X5Q, 2W4F). However, we did not see any ^15^N-^1^H chemical shift difference in the HSQC spectra of the amino acid residues comprising this region between the free and bound protein. This suggests that the extra C-terminal structure is likely present in the free protein too, and might function as an extra scaffold for stability. From the Scribble PDZ1 HSQC titrations spectra, we observed chemical shift perturbations for both the RPS6KA2 and MCC peptides (phosphorylated and unphosphorylated), further indicating that all four peptides interact with the Scribble PDZ1. The strength of the interactions could not be determined from these experiments only two point titrations (free and bound) were performed. Nevertheless, it gave an atomic information of the interaction being probed (see below).

**Figure 5.**
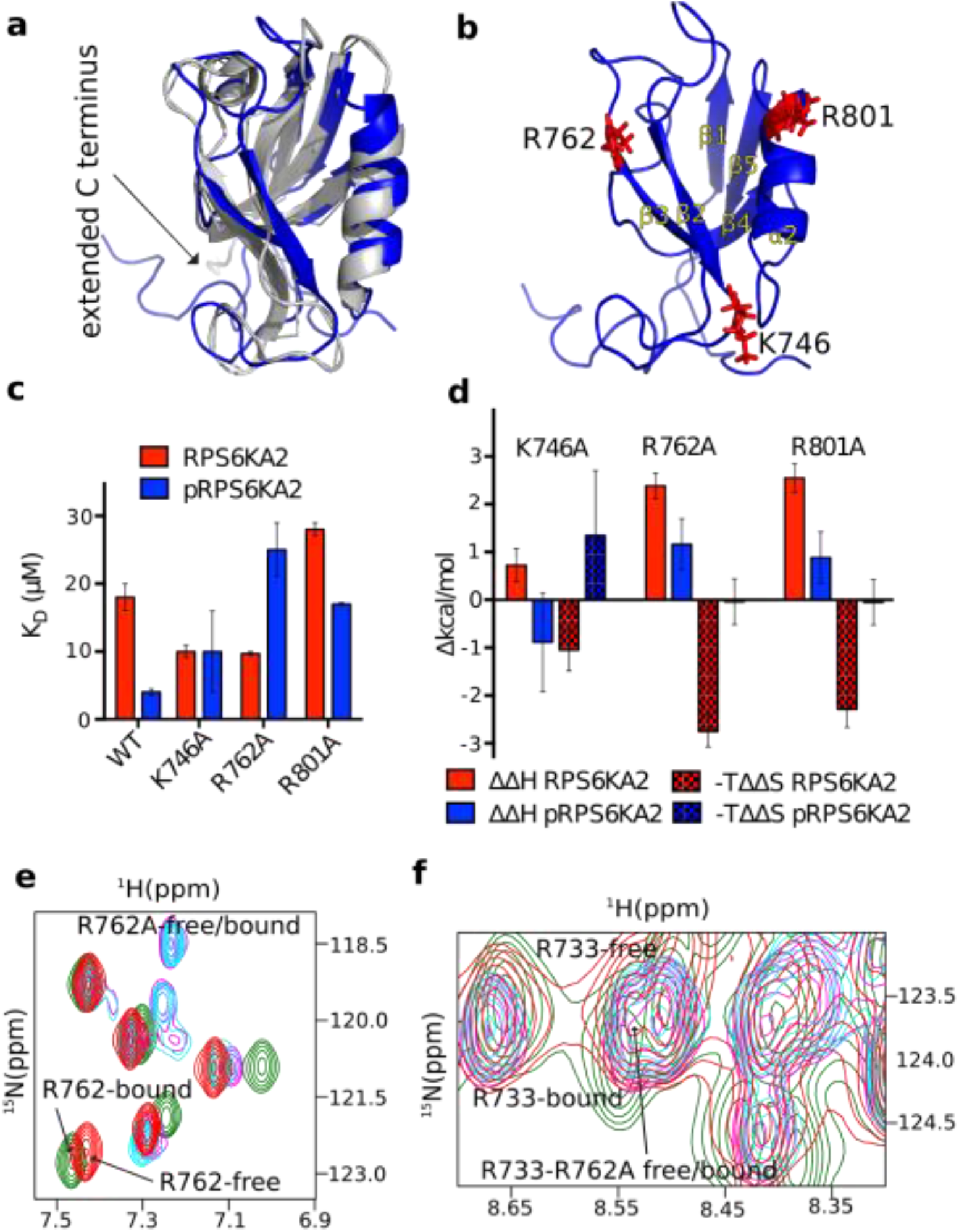
NMR structure of Scribble PDZ1 bound to phospho-RPS6KA2 and mutational analysis of specificity determining residues. **a)** Overlay of the previously published unliganded structure of Scribble PDZ1 (grey; PDB code 1x5g) and the NMR structure (blue; PDB code 6ESP) of the protein bound to MKRLTpSTRL-coo- (peptide not shown). **b)** Structure of the phosphopeptide-bound Scribble PDZ1 showing the positively charged residues around the binding pocket. **c)** Equilibrium binding constants for the binding between Scribble PDZ1 wild-type and mutants (K746A, R762A and R801A) and RPS6KA2 (unphosphorylated and p-3 phosphorylated) as determined through ITC titrations. **d)** Changes in the thermodynamic parameters upon mutation *(∆∆H*^*mutation*^ *and -T∆∆S*^*mutation*^) as determined through ITC titrations. **e)** Section of the ^15^N-^1^H HSQC spectra of free Scribble PDZ1 (red), bound to the phospho-RPS6KA2 peptide (green), R762A- mutant free (magenta) and R762A- mutant bound to phospho-RPS6KA2 (cyan). The R762 residue experience chemical shift perturbation only in the wild-type PDZ1. **f)** Slice of the ^15^N-^1^H HSQC showing the R733 residue for the respective proteins as in (e). R733 is critical for mediating the carboxylate at the c-terminus and its position does not change in the wild-type bound or the R762A bound showing that the carboxylate is still making the important interaction.

The NMR structure was determined with labeled protein and unlabeled, phosphorylated RPS6KA2 peptide (Table S2). As such only the protein was visible. However, we observed from the structure that K746, R762 and R801 are all surface-exposed and around the binding pocket (Fig. 5b, Fig S2). In addition, the ^15^N-^1^H titration experiments revealed that these residues experienced characteristic shift changes. We hypothesized that one or a combination of these positively charged residues are responsible for stabilizing the extra charges brought in by the phosphate group at position p-3. In order to test whether is the case we introduced single point mutations replacing the basic residues with neutral Ala (K746A, R762A and R801A) and performed ITC experiments as above for phosphorylated and unphosphorylated peptides. Each of the three substitutions conferred reduced affinities for the phosphopeptide, with the R762A mutation having the most pronounced effect on the affinity for phosphopeptide. The R801A mutation has also caused a reduced affinity for the unphosphorylated ligand. In contrast, the K746 and R762A mutations resulted in higher affinities for the unphosphorylated ligand; in case of the R762A mutation this resulted in a variant where phosphorylation has a disabling effect on ligand binding, and R762 can hence be considered a gate keeper residue providing Scribble PDZ1 with a specificity for p-3 phosphorylated RPS6KA2.

The effect of the R762A mutation on phosphopeptide binding is caused by a change in the enthalpic contribution to binding, and the residue also points into the cavity likely to be occupied by the phosphate group (Fig. S2). The effect of K746A is a bit unclear but perhaps it plays a role in steering the charged residues at −6 and −7 of the peptide. This observation comes from the HSQC titration experiment since both the phosphorylated and unphosphorylated peptides from MCC and RPS6KA2 all had huge chemical shift perturbation on K746, which is located at the end of the binding pocket. R801 on the other hand might play a steering role of both the C-terminal carboxylate and the phosphate group that are located in its vicinity.

To further explore the effect of the R762A mutation, we performed ^1^H-^15^N HSQC titrations of Scribble PDZ1 R762A and unphosphorylated and phosphorylated RPS6KA2. This titration revealed that the large conformational change exhibited by wild-type Scribble PDZ1 upon binding phospho-RPS6KA2 is lost in the R762A mutant (Fig. 5e). In contrast, the position of R733, which is critical for mediating interactions with the carboxylate at the C-terminus does not change in the wild-type bound or the R762A-bound forms, showing that the carboxylate of the peptide is still making the important interaction (Fig. 5f), supporting that the domain is still functional in binding unphosphorylated and phosphorylated RPS6KA2.

Our structural model, titration and mutagenesis analysis provide mechanistic insights into how Scribble PDZ1 bind the p-3 phosphorylated RPS6KA2 peptide and corroborate our results mentioned above.

### On the generality of PDZ domains and phosphopeptide interactions

The literature on ligands containing Ser/Thr phosphorylation sites that enable PDZ domain interactions is sparse. Nevertheless, we can still draw general conclusions about phosphopeptide-binding and PDZ domains. To this end we made a structure-based sequence alignment of the PDZ domains (this study) together with the previously suggested phosphopeptide binding PDZ domains of SNX27 and Shank1 (Fig. 6a). Through our study, we show that Scribble PDZ1 interacts with p-1 and p-3 phosphorylated ligands, and the phosphomimetic ProP-PD results suggest that Scribble PDZ2 and PDZ3 has similar preferences. For Scribble PDZ1 we found that binding of MKRLTpSTRL-coo- is connected to a conformational change of the binding pocket and relies on R762 as a key determinant for the selectivity for the p-3 phosphorylated ligand over the unphosphorylated ligands. A basic residue (Arg/Lys) is present at the same position in Scribble PDZ2 and PDZ3, suggesting that these domains may bind phosphopeptides in a similar fashion (Fig. 6). The presence of a basic residue in this position may thus serve as a first indication of phosphopeptide binding PDZ domains. Indeed, a Lys at the corresponding position was also previously implicated in the interaction between Tiam1 PDZ and the p-1 Tyr phosphorylated peptide of syndecan1^37^.

**Figure 6.**
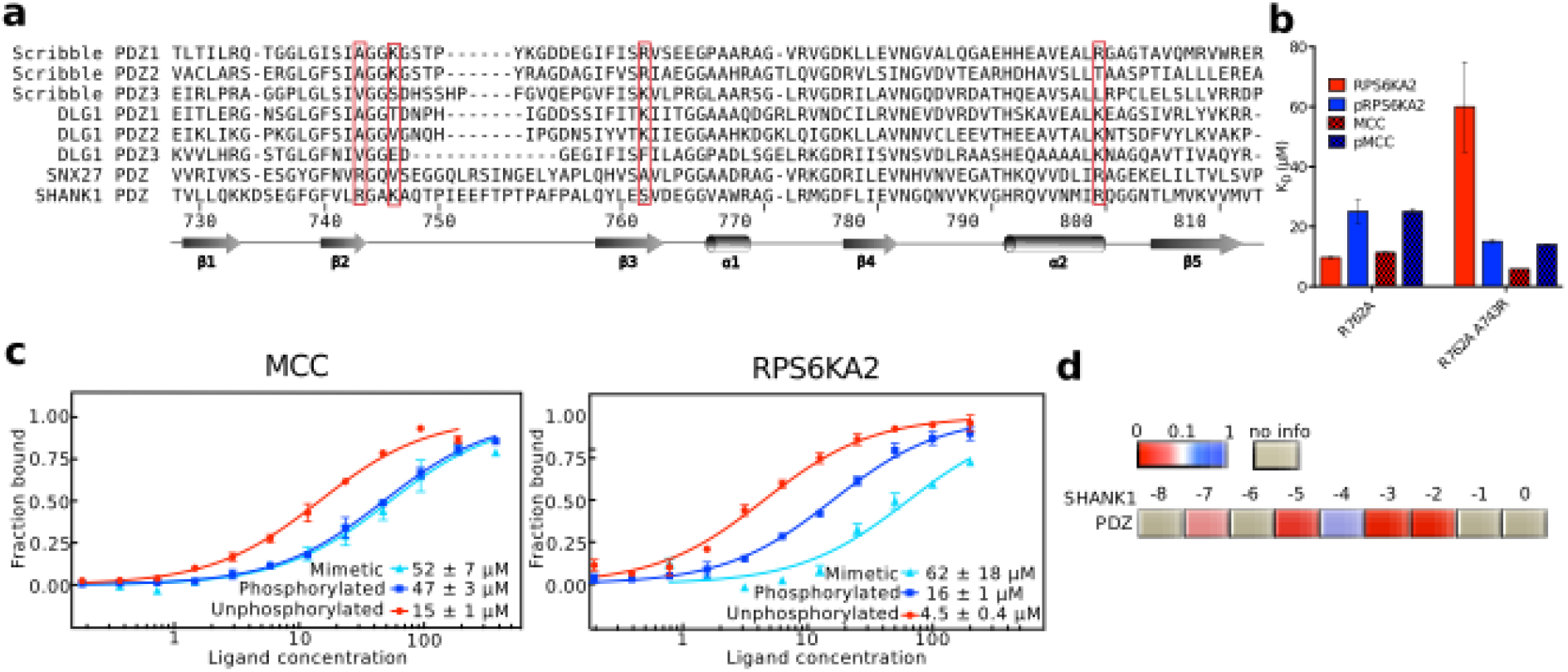
Plasticity of PDZ domain phosphopeptide shown by mutational analysis of Scribble R762A. The interactions of Shank1 PDZ domain were in contrast found to be disabled by ligand phosphorylation. a. Structure-based sequence alignment of selected PDZ domains. The amino acid numbering is based on full-length Scribble. b. Histogram of equilibrium dissociation constants for Scribble PDZ1 single (R726A) and double (R726A/A743R) mutants with unphosphorylated and phosphorylated peptides of MCC and RPS6KA2. c. MST affinity measurements using FITC-labelled peptides (WT, phosphorylated and phosphomimetic) of MCC and RPS6KA2. A fixed peptide concentration (50 nM) was titrated with varying concentration of Shank1 PDZ. *K*_*D*_ values were determined using thermophoresis + T-jump signal for data analysis (n=3 error bars represent SD). d. Scoring matrices of the effects of phosphomimetic mutations on the phosphomimetic ProP-PD results of Shank1 PDZ.

The PDZ domains of SNX27 and Shank1 lack a basic residue in the position corresponding to Scribble R762, but instead both have an Arg in β2 (corresponding to Scribble A743) that directly lines the peptide binding pocket. During the course of this study, the structures of SNX27 PDZ bound to phosphorylated peptides (p-3, p-5 and p-6) were reported, which revealed that SNX27 PDZ stabilizes the phosphate group through a network of residues including the Arg in β2^24^. The SNX27 PDZ domain and Scribble PDZ1 thus accomplish p-3 phosphorylated ligand binding in distinct manners. To explore the plasticity of the phosphopeptide binding, we introduced an Arg in β2 (A743R) in the Scribble PDZ1 R762A background. We determined the affinities for the unphosphoylated and phosphorylated MCC (p-1) and RPS6KA2 (p-3) peptides through ITC (Fig. 6b; Table S3). We found that the R762A/A743R mutation did not confer any specificity for the p-1 phosphorylated MCC peptide but it restored a preference for p-3 phosphorylated RPS6KA2. Thus, the specificity of Scribble PDZ1 for phosphopeptides can easily be tuned, and it is likely that this plasticity for phosphopeptide binding is shared with several other members of the PDZ domain family.

Finally, as noted above, Shank1 PDZ was previously suggested to bind both phosphorylated and unphosphorylated RPS6KA2 based on Y2H experiments with wild-type and phosphomimetic RPS6KA2 constructs^38^. To confirm the phosphopeptide binding of Shank1 PDZ we determined its affinities for the unphosphorylated and phosphorylated MCC and RPS6KA2 peptides through MST experiments. We found that Shank1 PDZ bind all peptides but that phosphorylation has a disabling effect (3x for MCC, 3.6x for RPS6KA2) (Fig. 6c, Table S3). To further explore the disabling of Shank1 PDZ binding upon ligand phosphorylation, we used the protein as a bait in selection against our phosphomimetic ProP-PD resource. We identified 11 protein ligands, including the wild-type but not phosphomimetic RPS6KA2 consistent with the MST data (Table S4). We further found that Shank1 PDZ interactions appear to be generally disabled by phosphomimetic mutations (Fig 6d), suggesting that although Shank1 PDZ and the PDZ domains of Scribble may have partially overlapping ligands, the interactions are likely differentially regulated by ligand phosphorylation.

### Discussion

We created a phage display library comprising all C-terminal peptides in the human proteome with known or putative Ser/Thr phosphorylation sites, and phosphomimetic variants thereof. We demonstrated the power of using such a phosphomimetic ProP-PD library together with NGS to identify interactions of potential biological relevance that can be enabled and disabled by Ser/Thr phosphorylation. The focus of this study is on the C-terminal regions of the proteome, and interactions with PDZ domains, but it can be envisioned to create such libraries that cover a large proportion of the Ser/Thr phosphosites in the disordered regions of the human proteome. Further studies with more extensive phosphomimetic ProP-PD libraries and alternative sets of phosphopeptide binding domains will establish the general applicability of the approach. We expect that the approach will prove to be feasible for probing phospho-regulated interactions on large-scale, in particular for domains that have an inherent affinity for their unphosphorylated ligands that is enhanced by the addition of the extra charges. The approach also offers an efficient way to pinpoint interactions that are negatively regulated by ligand phosphorylation. The method is likely less suited for proteins with strict requirement on the phosphate group for binding. To explore such cases, peptide SPOT arrays or protein arrays probed with phosphopeptides would be more appropriate experimental methods. In an expanded form, phosphomimetic ProP-PD can be a highly useful tool for assigning functional roles to the phosphoproteome, and for linking of phosphorylation switched SLiMs to their respective binding domains, thereby contributing to solving the phosphorylation code.

From the perspective of PDZ domains, we provide evidences for a dynamic switch-like mechanism, where phosphorylation at p-1 or p-3 enable interactions with Scribble PDZ domains, while disabling interactions with other PDZ domains, such as Shank1 PDZ. Such effects are likely to be of functional importance in a cellular complex where the composition of PDZ protein complexes are determined by the dynamic equilibriums between the relative affinities of the interacting proteins and the relative protein concentrations.

From a structural point, we found that Scribble PDZ1 binding of a p-3 phosphorylated ligand was associated with a conformational change of the peptide binding pocket, which allows the phosphoryl group of the peptide to interact with an Arg in β3. Utilizing a basic residue at this position for stabilizing interactions with a negatively charged phosphoryl group may be a rather general strategy among PDZ domains. Indeed, the corresponding Arg in the Tiam1 PDZ scaffold was previously found to be crucial for binding a p-1 Tyr phosphorylated syntenin1 peptide although no major conformational changes of the protein was associated with this interaction ^37^. Binding of p-3 phosphorylated ligands can also involve an Arg in β2, as recently shown for the SNX27 PDZ domain, and here confirmed through mutational analysis of Scribble PDZ1. However, as shown for Shank1 PDZ, an Arg in β2 is not always sufficient to provide specificity for phosphorylated ligands. A basic residue in the key positions in β3 or in β2 may thus be required but not sufficient to support phosphopeptide binding. Further dedicated studies will be required before we can provide a comprehensive picture of how ligand phosphorylation tunes interactions with different PDZ domain family members, and how this contributes to the diversify and fine-tune the specificities across the PDZ domain family. Based on our results, we expect that phosphorylation switches of PDZ domain interactions will be more common than previously appreciated. Phosphomimetic ProP-PD will be a powerful approach in the continued quest for elucidating the phosphorylation switching of PDZ domain mediated interactions as well as of other SLiM-based interactions.

## Acknowledgements

This work was supported by grants to from the Swedish research council (YI, 2012-05092 and 2016-04965), the Wiberg foundation (YI), the Carl Trygger foundation (YI, CTS15:226), CNC was supported by a Wenner-Gren return fellow program starting grant. The phagemid used for library construction was generously provided by the Sidhu lab (University of Toronto), together with expression constructs encoding GST-tagged proteins. The plasmid encoding GFP-tagged full-length human Scribble was generously provided by JP Borg (INSERM Marseille).

Author contributions: GN, AR, CNC and YI conceived experiments. AR designed the phage library, GN purified proteins, created the phage library, performed biophysical and cell based validations. CNC determined the NMR structure. MA performed affinity determinations. JO performed peptide NMR titrations. PG provided the CYANA program and participated in structure calculation. GN, AR, CNC and YI wrote the manuscript.

## SUPPLEMENTAL INFORMATION

### Supplemental Methods

**Figure S1.** Microscale thermophoresis affinity measurements of FITC-labeled peptides and recombinant PDZ domains.

**Figure S2**. Supplementary NMR figures of Scribble PDZ1.

**Table S1.** High-confidence sets of annotated ligands of the PDZ domains of Scribble and DLG1 generated through phosphomimetic ProP-PD.

**Table S2**. NMR statistics.

**Table S3**. Summary of dissociation constants.

**Table S4**. Annotated ligands of Shank1 PDZ suggested by phosphomimetic ProP-PD.

